# Marine water environmental DNA metabarcoding provides a comprehensive fish diversity assessment and reveals spatial patterns in a large oceanic area

**DOI:** 10.1101/864710

**Authors:** Natalia Fraija-Fernández, Marie-Catherine Bouquieaux, Anaïs Rey, Iñaki Mendibil, Unai Cotano, Xabier Irigoien, María Santos, Naiara Rodríguez-Ezpeleta

## Abstract

Current methods for monitoring marine fish diversity mostly rely on trawling surveys, which are invasive, costly and time-consuming. Moreover, these methods are selective, targeting a subset of species at the time, and can be inaccessible to certain areas. Here, we explore the potential of environmental DNA (eDNA), the DNA present in the water column as part of shed cells, tissues or mucus, to provide comprehensive information about fish diversity in a large marine area. Further, eDNA results were compared to the fish diversity obtained in pelagic trawls. A total of 44 5L-water samples were collected onboard a wide-scale oceanographic survey covering about 120,000 square kilometres in Northeast Atlantic Ocean. A short region of the 12S rRNA gene was amplified and sequenced through metabarcoding generating almost 3.5 million quality-filtered reads. Trawl and eDNA samples resulted in the same most abundant species (European anchovy, European pilchard, Atlantic mackerel and blue whiting), but eDNA metabarcoding resulted in more detected fish and elasmobranch species (116) than trawling (16). Although an overall correlation between fish biomass and number of reads was observed, some species deviated from the common trend, which could be explained by inherent biases of each of the methods. Species distribution patterns inferred from eDNA metabarcoding data coincided with current ecological knowledge of the species, suggesting that eDNA has the potential to draw sound ecological conclusions that can contribute to fish surveillance programs. Our results support eDNA metabarcoding for broad scale marine fish diversity monitoring in the context of Directives such as the Common Fisheries Policy or the Marine Strategy Framework Directive.

## Introduction

Monitoring of marine biodiversity provides a baseline for policy implementation towards a sustainable use of the marine environment and its resources. Among the traditional methods for surveying marine fauna, trawling has been widely used, as identification and quantification of large volumes of organisms is considered a reliable method for monitoring fish and other marine animal populations (ICES, 2015; Massé, Uriarte, Angélico, & Carrera, 2018). Fish surveys using trawls are conditioned by the gear’s own characteristics (e.g., mesh size, area of opening) and deployment parameters (e.g., towing speed, depth and diel variation) (Heino et al., 2011). Consequently, besides being invasive and time consuming, fish trawling in pelagic environments can be largely selective affecting diversity estimates and knowledge of species composition (Fraser, Greenstreet, & Piet, 2007; ICES, 2004). For instance, due to their large body size, fast swimming speed, and in some cases, scarcity, many elasmobranch species are not thoroughly surveyed (Rago, 2004). Therefore, alternative methods are needed, and advances in DNA sequencing and bioinformatics have opened new avenues to assess marine biodiversity in a non-invasive manner (Danovaro et al., 2016; Rees, Maddison, Middleditch, Patmore, & Gough, 2014).

In particular, the analysis of environmental DNA (eDNA), that is, the genetic material shed and excreted by organisms to the environment, to characterise the biological communities present in an environment (P. Taberlet, Coissac, Hajibabaei, & Rieseberg, 2012) is gaining increasing attention for monitoring aquatic environments (Thomsen & Willerslev, 2015). Community composition can be inferred from eDNA samples through metabarcoding, whereby the eDNA is collected from the water column through filtering, selectively amplified through PCR using primers targeting a given barcode from a particular taxonomic group and sequenced (Pierre Taberlet, Coissac, Pompanon, Brochmann, & Willerslev, 2012). The resulting sequences are then compared against a reference database to perform biodiversity inventories (Deiner et al., 2017). Most studies using eDNA metabarcoding for monitoring fish communities are based in freshwater environments and have shown that eDNA metabarcoding provides overall estimates that are equivalent or superior to traditional methods such as visual surveys, trawling or electrofishing (Hänfling et al., 2016; Minamoto, Yamanaka, Takahara, Honjo, & Kawabata, 2012; Pont et al., 2018).

As opposed to freshwater systems, the marine environment has in general a larger water-volume to fish biomass ratio and is influenced by currents, implying that the eDNA is less concentrated and disperses quicker (Hansen, Bekkevold, Clausen, & Nielsen, 2018). This, coupled with a higher sympatric marine fish diversity, suggests that monitoring fish diversity through eDNA sampling could be particularly challenging in the marine environment. Indeed, only a handful of studies have applied eDNA metabarcoding for monitoring fish and elasmobranchs in natural marine environments (e.g., O’Donnell et al., 2017; Stat et al., 2017). Among them, only a few have compared eDNA and other traditional surveying methods and are based on a very small area of a few square kilometres either in ports (Jeunen et al., 2019; Sigsgaard et al., 2017; Thomsen et al., 2012) or in coastal areas (Andruszkiewicz et al., 2017; DiBattista et al., 2017; Yamamoto et al., 2017) or have performed comparisons at family level taxonomic assignments (Thomsen et al., 2016). Thus, although these studies envision eDNA metabarcoding as a promising method for non-invasive, faster, more efficient and reliable marine surveys, this needs still to be tested in the context of a fishery survey covering a broad marine area.

The Bay of Biscay is a biogeographical area in the North Atlantic Region covering more than 220,000 Km^2^, at which the main economic activities include commercial fishing. Large populations of species such as the European anchovy *Engraulis encrasicolus*, the European pilchard *Sardina pilchardus*, the European hake *Merluccius merluccius*, the Atlantic Mackerel *Scomber scombrus* and the Atlantic horse mackerel *Trachurus trachurus* are dominant in the area (ICES, 2018). Fish diversity in the Bay of Biscay has been accounted using mainly observational methods, fish trawling and acoustic surveys; thus, there is room for incorporating and assessing the performance of eDNA-based surveys. This paper aims to test the potential of eDNA metabarcoding to assess the fish community composition in a large marine area, such as the Bay of Biscay. For that aim, we have compared eDNA metabarcoding based estimates with those derived from fishing trawls catches and have related eDNA metabarcoding based estimates with the known spatial distribution and ecological patterns of the species in the area.

## Methods

### Sample collection

Fish catches and water samples were collected during the BIOMAN 2017 survey (Santos, Ibaibarriaga, Louzao, Korta, & Uriarte, 2018) between the 5^th^ and the 29^th^ of May 2017 covering the area of about 120,000 km^2^ between the French continental shelf and the Spanish shelf (Figure 1) on board the Emma Bardán and Ramón Margalef research vessels. Fish catches were obtained on board the R/V Emma Bardán pelagic trawler. The trawl had an 8 mm mesh size cod end, and towing time and speed were 40 min and 4 knots, respectively. A total of 44 stations were used for trawling. Although station depths varied between 26 and 3000 m, the maximum fishing depth was 156 m. Onboard, fish were morphologically identified to species level or, when doubt, to the smallest taxonomic rank (e.g., family or genus). Biomass estimates were standardised as Kg caught per taxa. Water samples were collected on board the R/V Ramón Margalef research vessel using the continuous circuit intake of the ship at 4.4 m depth in 44 additional stations (Figure 1). A total of 5 L sea water per station was filtered through Sterivex 0.45 µm pore size enclosed filters (Millipore) with a peristaltic pump and kept at −20 °C until further processing.

**Figure 1.**
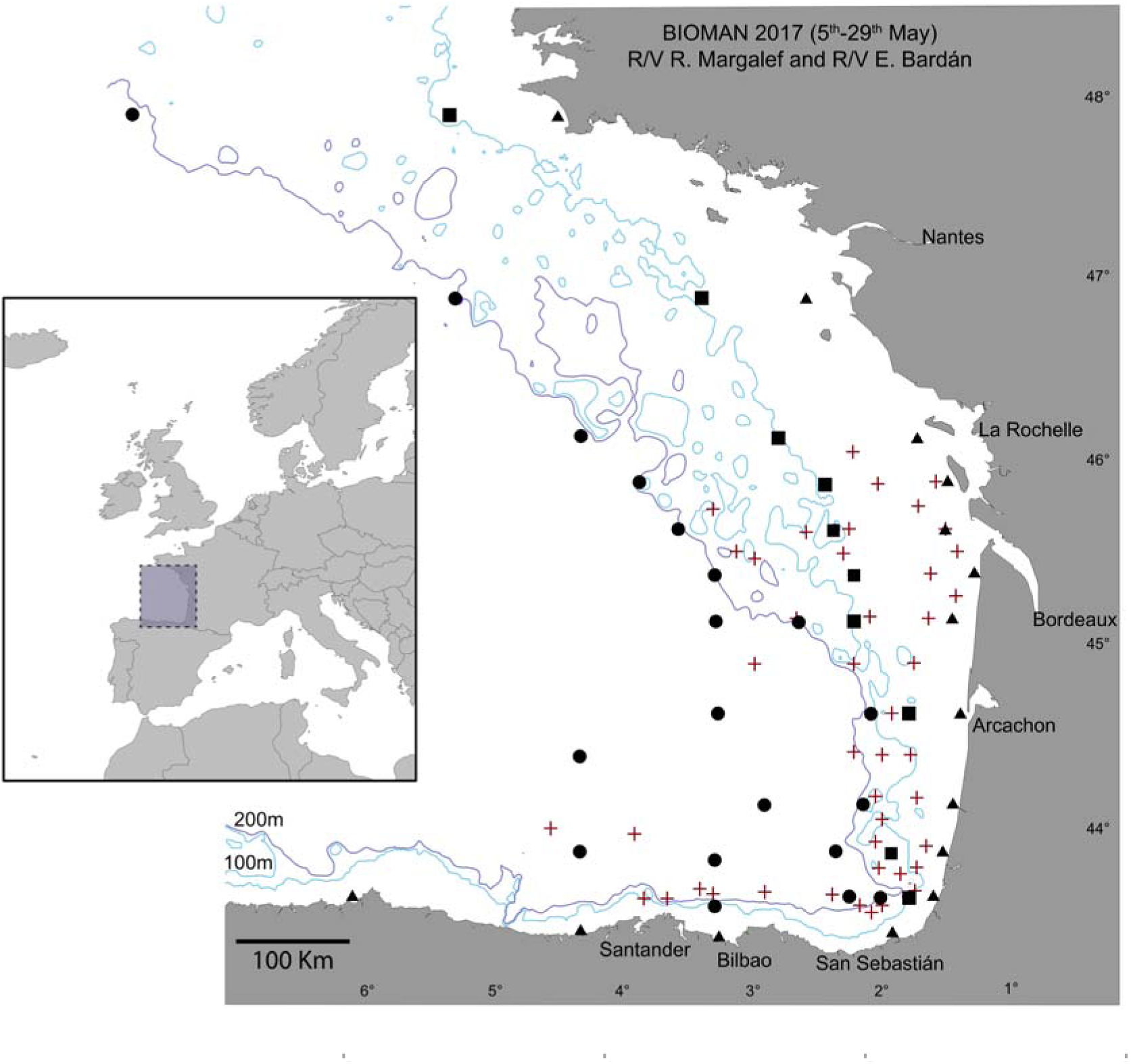
Study area and sampling sites for the BIOMAN 2017 survey in the Bay of Biscay. Triangles represent eDNA sampling sites where station depth was <90 m, squares, eDNA sampling sites with depths between 90 m and 127 m, and circles, eDNA sampling sites with >127 m depths. Crosses are located where pelagic fishing trawls were deployed. 100 m and 200 m isobaths are shown.

### DNA extraction and amplicon library preparation

DNA extractions were performed using the DNeasy® blood and tissue kit (Qiagen) following the modified protocol for DNA extraction from Sterivex filters without preservation buffer by Spens et al. (2017). DNA concentration was measured with the Quant-iT dsDNA HS assay kit using a Qubit® 2.0 Fluorometer (Life Technologies, California, USA). DNA from all 44 samples was amplified with the teleo_F/telo_R primer pair (hereafter ‘teleo’), targeting a short region (□60 bp) of the mitochondrial 12S rRNA gene, combined with the human blocking primer teleo_blk (Valentini et al., 2016). PCR amplifications were done in a final volume of 20 µl including 10 µl of 2X Phusion Master Mix (ThermoScientific, Massachusetss, USA), 0.4 µl of each amplification primer (final concentration of 0.2 µM), 4 µl of teleo_blk (final concentration of 2 µM), 3.2 µl of MilliQ water and 2 µl of 10 ng/µl template DNA. Samples from 4 stations were also amplified i) using the same procedure but without the blocking primer and ii) using the mlCOIintF/dgHCO2198 primer pair (hereafter ‘mlCOI’), targeting a short region (□310 bp) of the COI gene (Meyer, 2003). The thermocycling profile for PCR amplification included 3 min at 98 °C; 40 or 35 cycles (for ‘teleo’ and ‘mlCOI’, respectively) of 10 s at 98 °C, 30 s at 55 or 46 °C (for ‘teleo’ and ‘mlCOI’, respectively) and 45 s at 72 °C, and finally, 10 min at 72 °C. PCR products were purified using AMPure XP beads (Beckman Coulter, California, USA) following manufacturer’s instructions and used as templates for the generation of 12 x 8 dual-indexed amplicons in the second PCR reaction following the ‘16S Metagenomic Sequence Library Preparation’ protocol (Illumina, California, USA) using the Nextera XT Index Kit (Illumina, California, USA). Multiplexed PCR products were purified using the AMPure XP beads, quantified using Quant-iT dsDNA HS assay kit using a Qubit® 2.0 Fluorometer (Life Technologies, California, USA) and adjusted to 4 nM. 5 µl of each sample were pooled, checked for size and concentration using the Agilent 2100 bioanalyzer (Agilent Technologies, California, USA), sequenced using the 2 x 300 paired end protocol on the Illumina MiSeq platform (Illumina, California, USA) and demultiplexed based on their barcode sequences.

### Reference database

Two reference databases were created for the ‘teleo’ barcode. A first ‘global’ database included all Chordata 12S rRNA and complete mitochondrial genome sequences available from GenBank (accessed in February 2018). By performing an all-against-all BLAST (Altschul, Gish, Miller, Myers, & Lipman, 1990), potential sources of contamination or erroneous taxonomic assignments were removed such as human contaminations (e.g., non-human labelled sequences that matched at 100% identity with the *Homo sapiens* 12S rRNA sequence) or cross-contaminated sequences (e.g., sequences arising from the same study that, even when belonging to different genus, were 100% identical). All sequences were trimmed to the ‘teleo’ region. Taxonomy for the GenBank sequences was retrieved using E-utilties (Sayers, 2008) and modified to match that of the World Register of Marine Species: WoRMS (Horton et al., 2018), forcing for seven taxonomic levels, i.e., Phylum, Subphylum, Class, Order, Family, Genus, Species. This ‘global’ reference database contains 10,284 ‘teleo’ region sequences. For the second database, only sequences from target species were retrieved so that more exhaustive error checking was possible. The list of the 1,858 fish species expected in the Northeast Atlantic and Mediterranean areas was compiled from FishBase (http://www.fishbase.org) and their corresponding scientific names and sequences were obtained from NCBI (https://www.ncbi.nlm.nih.gov). For the retrieved records, only those covering the ‘teleo’ region were selected and aligned (Table S1). A phylogenetic tree was built with RAxML (Stamatakis, 2014) using the GTR-CAT model and visualized with iTOl (Letunic & Bork, 2016). The tree was visually inspected and the records corresponding to misplaced species were removed from the database. This ‘local’ reference database contains ‘teleo’ region sequences of 612 species. For the ‘mlCOI’ barcode, the reference database consisted in the COI sequences and their corresponding taxonomy obtained from the BOLD (Ratnasingham & Hebert, 2007) database.

### Read pre-processing, clustering and taxonomic assignment

Quality of raw demultiplexed reads was verified with *FASTQC* (Andrews, 2010). Forward and reverse primers, were removed with *cutadapt* (Martin, 2011) allowing a maximum error rate of 20%, discarding read pairs that do not contain the two primer sequences and retaining only those reads longer than 30 nucleotides. Paired reads were merged using *pear* (Zhang, Kobert, Flouri, & Stamatakis, 2014) with a minimum overlap of 20 nucleotides. Pairs with average quality lower than 25 Phred score were removed using *Trimmomatic* (Bolger, Lohse, & Usadel, 2014). *mothur* (Schloss et al., 2009) was used to remove reads i) not covering the target region, ii) shorter than 40 or 313 nucleotides, for ‘teleo’ and ‘mlCOI’, respectively, iii) containing ambiguous positions and iv) being potential chimeras, which were detected based on the *UCHIME* algorithm (Edgar, Haas, Clemente, Quince, & Knight, 2011). Reads were clustered into OTUs using *vsearch* (Rognes, Flouri, Nichols, Quince, & Mahé, 2016) at 97% similarity threshold and into SWARMs using *Swarm* (Mahé, Rognes, Quince, de Vargas, & Dunthorn, 2014) with a d value of 1. In both cases, the *LULU* post-clustering algorithm (Frøslev et al., 2017) was applied with a minimum threshold of sequence similarity for considering any OTU/SWARM as an error of 97%. Taxonomic assignment of unique reads and of representative sequences for each OTU or SWARM was performed using the naïve Bayesian classifier method (Wang, Garrity, Tiedje, & Cole, 2007) implemented in *mothur* using the 12S rRNA and COI databases described above. Reads with the same taxonomic assignment were clustered into phylotypes.

### Biodiversity analyses

Analyses were performed with the R packages *Phyloseq* (McMurdie and Holmes 2013) and *Vegan* (Oksanen et al., 2019). Sampling stations were classified into three categories considering their depth and distance from the coast (see Map in Figure 1): shallow stations where maximum station depth was <90 m, medium stations, when depth ranged between 90 and 127 m and, deep stations where depth was >127 m. To assess differences in fish diversity across sampling zones (i.e., according to shallow, medium and deep stations) we calculated the Bray-Curtis dissimilarity index for relative abundance of species with the function *ordinate* using only phylotypes with more than 10 reads. These distances were then ordinated using a non-metric Multidimensional Scaling (NMDS) as implemented in *Phyloseq* and differences between stations were tested with PERMANOVA (1,000 permutations) using the function *adonis* within the R package *Vegan* previous testing for homogeneity of variance using the function *betadisper*. A linear model was used on species with more than 1,000 reads, to test for the effect of the abundance of reads (previously standardised according to the overall number of reads and stations per zone), and the distance from the coast. An overall correlation between the log-transformed values of the number of reads obtained and the biomass caught per species was explored with the Pearson correlation coefficient as implemented in R package *Stats*. For an even geographic distribution between water and fish sampling sites, a total of nine water sampling sites north La Rochelle were removed for the correlation analysis. In addition, in order to compare eDNA and trawling based estimates at a smaller scale, we created groups of stations so that this comparison was possible. For that aim, we combined the data from all eDNA and trawling stations within < 20nm of each eDNA station in what we call mega-stations. A total of 30 mega-stations resulted. A mantel test as implemented in the R package *ade4* (Dray & Dufour, 2007) was used to explore differences between geographic and Bray-Curtis distances of the mega-stations. The list of species commonly reported from the Bay of Biscay was obtained mainly from i) Basterretxea, Oyarzabal, and Artetxe (2012), ii) the AZTI’s database on fish bottom trawling discards in the area gathered according to EU regulation 2017/1004 of 17 May 2017, iii) the data obtained from fish pelagic trawling during BIOMAN surveys from 2003 until 2019, iv) the ICES database for International Bottom Trawling Surveys available from www.ices.dk and v) the 2017 Pélagiques Gascogne (PELGAS) integrated survey (Mathieu et al., 2019).

## Results

### Data quality and overall taxonomic composition

We obtained a total of 4,640,913 raw ‘teleo’ reads from which 3,366,264 (72%) were retained after quality check for downstream analyses. The average number of ‘teleo’ reads per sample was 70,131 (Table S2). Using the ‘global’ database, 99.88% of the reads were classified as Actinopterygii or Elasmobranchii. The remaining 0.12% (4,343 reads) were classified as mammals (40.16%) and birds (9.60%), with half of the reads (50.24%) not classified into Class level (Figure S1). Only 14 reads in eight samples were specifically assigned to *H. sapiens*. From these, two samples did not include the specific blocking primer used, suggesting that samples held very little contamination from external sources. Using the ‘local’ database, 99.98% of the reads were classified either as Actinopterygii or Elasmobranchii and, depending on the clustering method used, the number of taxa recovered varied. SWARM clustering yielded 90 OTUs identified at the species level (including 95.5% of the reads) and vsearch, 109 (including 95% of the reads), whereas not clustering reads into OTUs, but using phylotypes, resulted in 116 Actinopterygii and Elasmobranchii species (including 95% of the reads) identified. Further analyses were based on phylotypes assigned to the species level (Table S3) as no additional information is provided by using OTU clustered reads. From the 116 identified species, 50 included more than 10 reads.

More than half of the reads are assigned to European anchovy, *E. encrasicolus* (51.67%), followed by European pilchard, *S. pilchardus* (27.67%), Atlantic mackerel, *Scomber scombrus* (4.96%), blue whiting, *Micromesistius poutassou* (2.36%), white seabream, *Diplodus sargus* (1.52%) and axillary seabream *Pagellus acarne* (1.20%), which together represent 89.4% of the reads (Figure 2A). A small percentage of the reads (0.27%) were classified as Elasmobranchii, including seven species such as the Greenland shark, *Somniosus microcephalus*, the blue shark, *Prionacea glauca* and the undulate ray, *Raja undulata* (Figure 2B).

**Figure 2.**
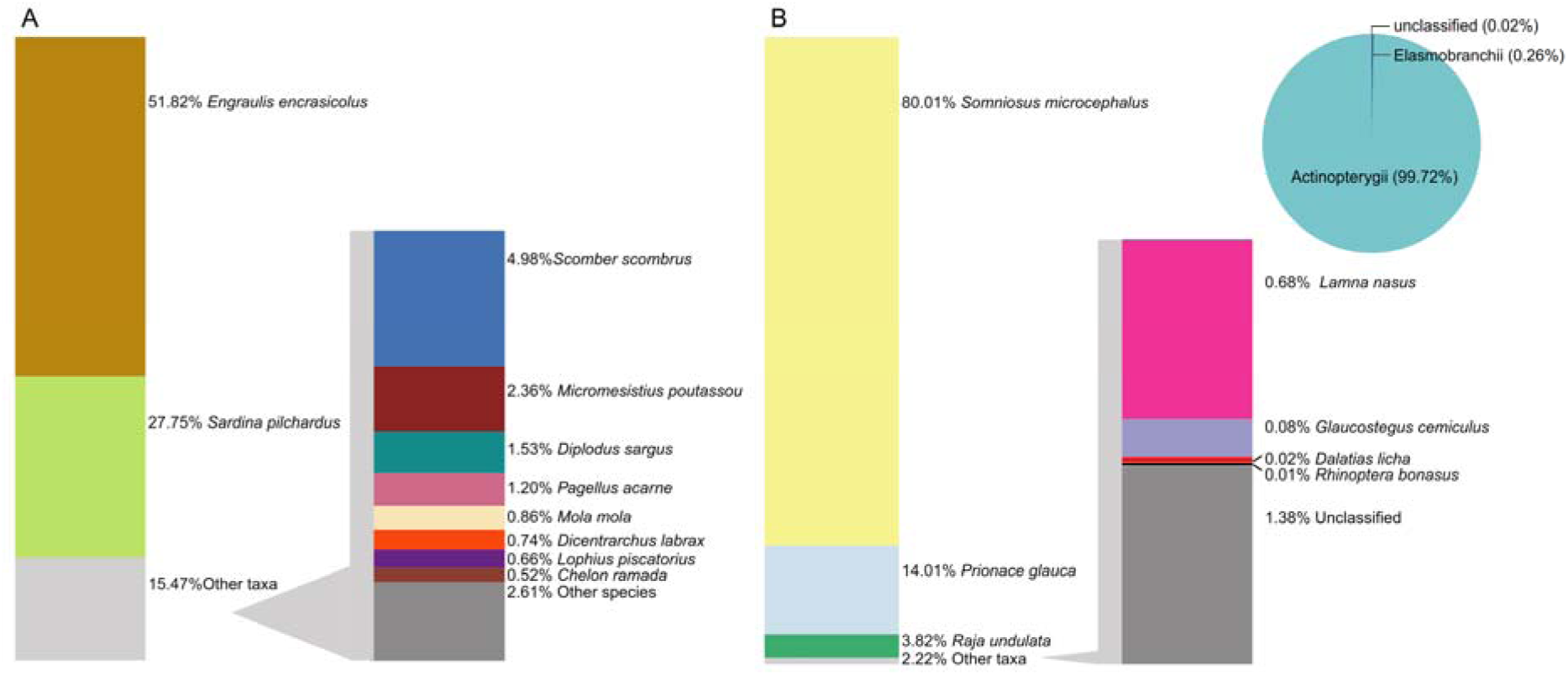
Relative number of reads (%) assigned to **(A)** Actinopterygii and **(B)** Elasmobranchii species recovered from eDNA metabarcoding. Note that 4.96% Actinopterygii were not classified into species level.

As for the four samples amplified with ‘mlCOI’ primers, we obtained 389,665 raw reads from which, 324,731 (83%) were retained for downstream analyses. The average number of ‘mlCOI’ reads per sample retained after quality filtering is 81,183 (Table S2). Using the BOLD database, 89.86% of the reads were classified into Phylum, 80.87% of which were metazoans, and among them 47.88% were classified as arthropods and 2.51% as chordates (Figure S2). Within chordates 74.56% of the reads were classified as Actinopterygii (1.87% of the overall reads), resulting in only seven taxa classified into species (Figure S2).

### Comparison with fish trawling

Trawling operations during the BIOMAN survey resulted in a total of 18 taxa caught, from which lanternfishes (Fam. Myctophidae) and mullets (*Mugil* sp.) were the only ones not classified into species level. Qualitatively, a total of 10 species were identified both from the eDNA and trawling catches (Figure 3A). Six species were collected during catches and not detected through eDNA, namely *Sprattus sprattus*, *Trachurus mediterraneus*, *Boops boops*, *Zeus faber*, *Trisopterus luscus*, and *Capros aper* (Table S4); from these, there are no sequences for *T. mediterraneus* and *B. boops* in the reference database and the fact that we find *T. minutus* in eDNA suggest that this could be actually *T. luscus*. To assess the relationship between the biomass of fish caught and the number of reads obtained through eDNA, data from *T. mediterraneus* and *T. trachurus* were combined into *Trachurus* spp. and that from *T. luscus* and *T. minutus* into *Trisopterus* spp. There was an overall correlation between fish biomass and number of reads per species although not significant at p < 0.05 (Figure 3B). *E. encrasicolus* was the most abundant species for both methods, whilst the relative abundance for some species like *Dicentrarchus labrax, M. poutassou* and *S. pilchardus* was higher when using eDNA. In contrast, the relative abundance of *M. merluccius, S. scombrus* and *Trachurus* spp. was higher in catches than when using eDNA (Figure 3B; Table S4). At a local scale, no significant correlation between eDNA and trawling based abundances was found (Mantel test, r = −0.04 p = 0.646). In fact, eDNA data showed a more constant abundance of the three most abundant species (*E. encrasicolus*, *S. pilchardus* and *S. scombrus*), compared to trawl data, which showed in general a higher number of species per station, except for those eight stations were *E. encrasicolus* was dominant (> 94% of the catch) (Figure S3).

**Figure 3.**
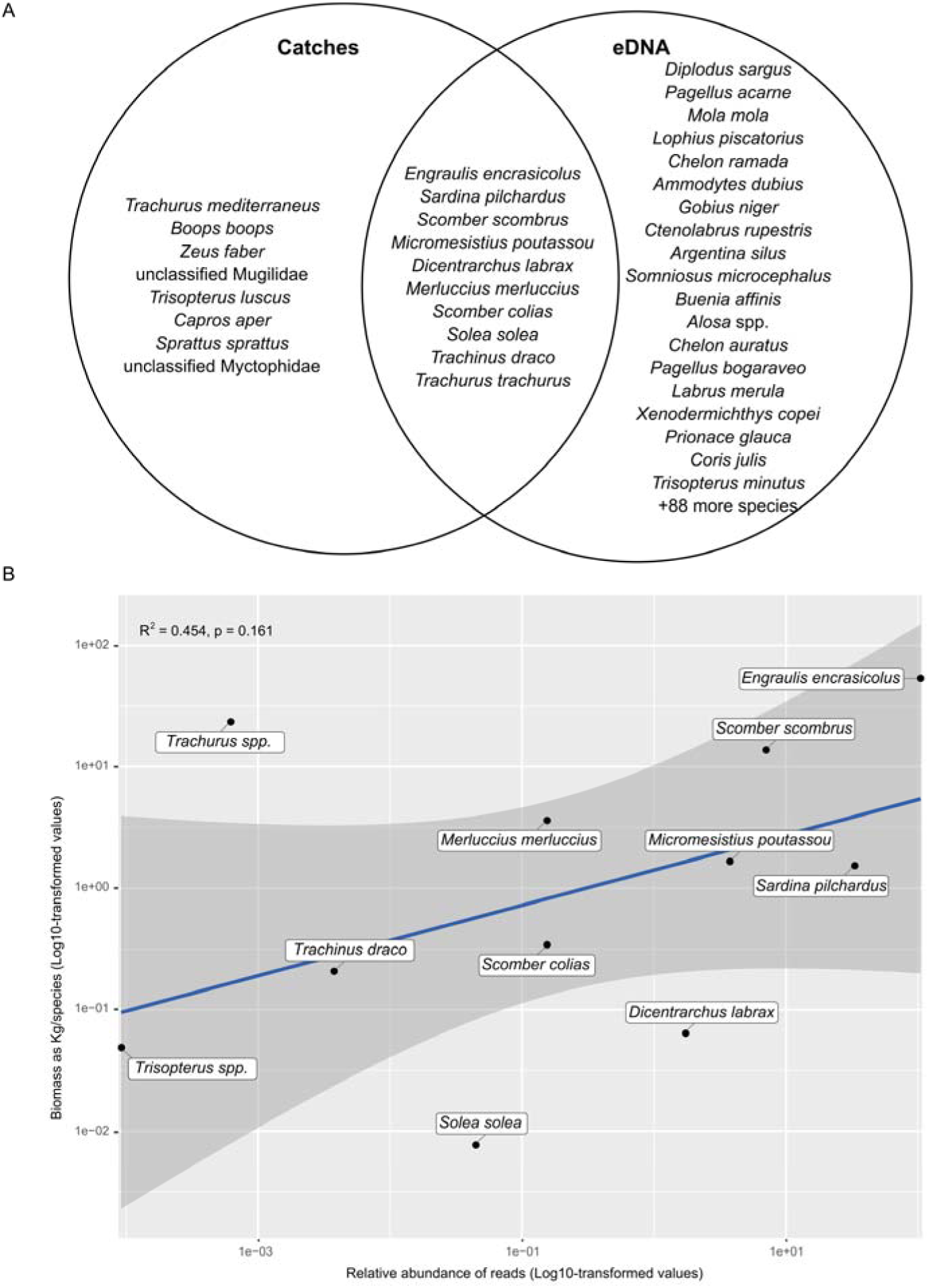
**(A)** Venn diagram showing fish species caught in trawls and detected through eDNA metabarcoding organised in decreasing order according to biomass or number of reads. **(B)** Relationship between the log10-transformed values for the number of reads and biomass in Kg from all fish species simultaneously found through eDNA and caught during fish trawling. Shaded area represents the 95% confidence interval of the linear regression.

### Species distribution patterns

A significant correlation was detected between species abundance differences and geographic distances between stations both for eDNA (R^2^ = 0.38 p < 0.01) and trawling stations (R^2^ = 0.20 p < 0.01). Yet, although for eDNA, pairs of distant stations tend to be more different in taxonomic composition, pairs of nearby stations cover the full range of Bray-Curtis distances (Figure 4). Comparisons between samples within same or distinct depth category (shallow, medium, deep) or within same or distinct sampling methods (eDNA, trawling) had no effect over the observed patterns (Figure S4).

**Figure 4.**
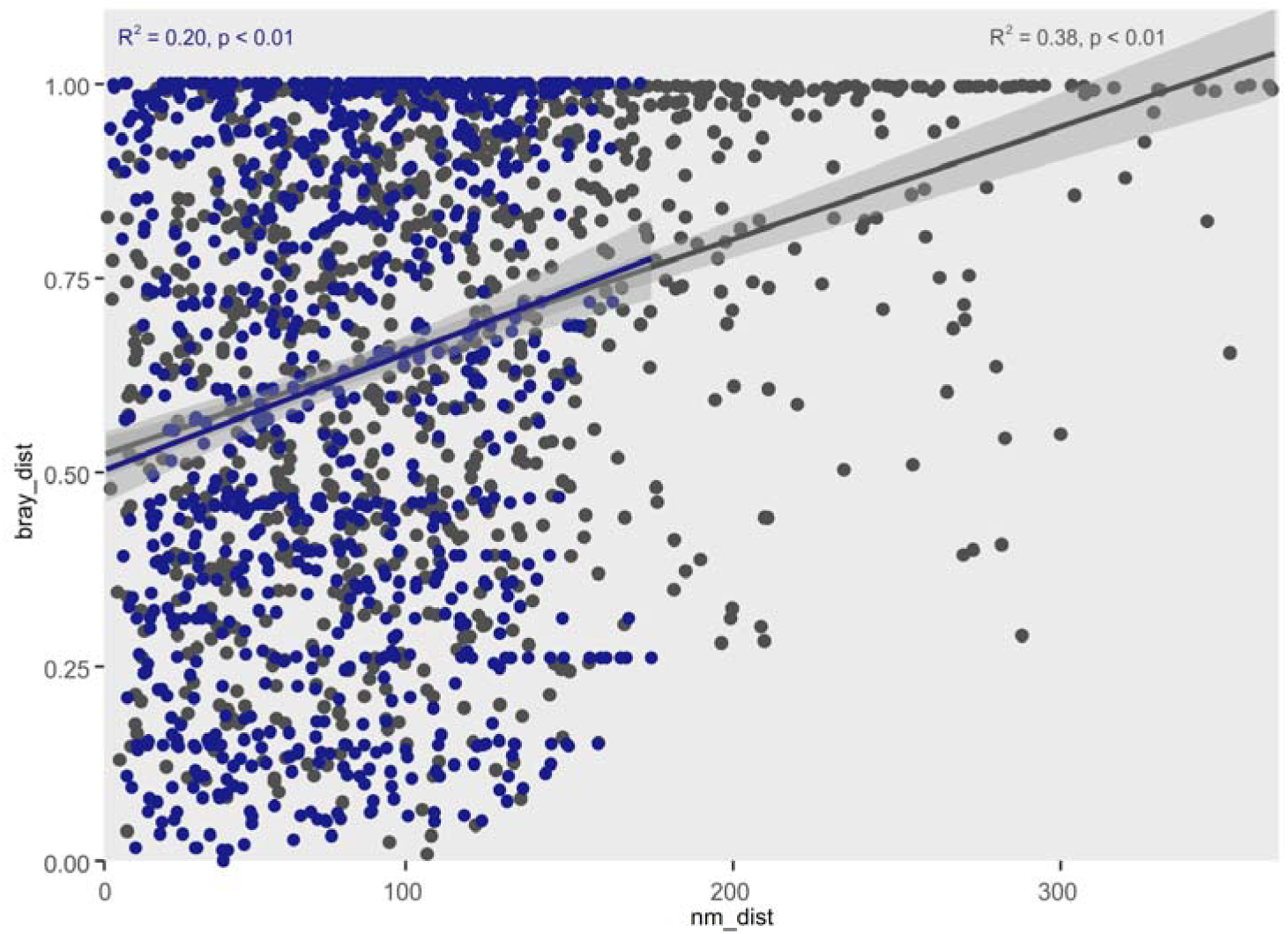
Scatterplot showing the overall relationship between Bray-Curtis distance and geographic distance between pairs of eDNA (grey) and trawling (blue) stations. Pearson correlation is shown for each data group. Shaded area represents the 95% confidence interval of the linear regression.

The overall compositional pattern of our data showed significant differences between species occurrence and sampling sites according to their zone (e.g., shallow, medium and deep stations) (PERMANOVA F_2,43_ = 2.24, *p* < 0.05) (Figure 5). Within the main species contributing to the spatial ordination of our data, two main groups can be broadly observed. On one side, species like *E. encrasicolus, M. merluccius, Coris julis S. scombrus, M. poutassou, Lophius piscatorius, S. microcephalus, Xenodermichthys copei*, and *P. glauca* tended to be more abundant in deeper stations and their relative abundances increased in sites >127m-deep (Figure 6). In contrast, a second loop in the spatial ordination of the data include other species such as *Gobius niger, Ammodytes dubius, D. sargus, Argentina silus, D. labrax, S. pilchardus, Mola mola* and *Scomber colias* (Figure 5). This information correlates with a pattern of higher abundance in <90m-deep sites for, e.g., *S. pilchardus, D. sargus, M. mola, A. dubius*, *D. labrax* and *S. colias* (Figure 4B). Relatively to the abundance of reads and station depth, four species, namely *A. silus*, *Glaucostegus cemiculus*, *G. niger* and *Pagellus bogaraveo*, remain unchanged between shallow and deep stations. Specifically, for elasmobranch species, a pattern nicely correlated with higher relative abundances of typical demersal species like *R. undulata* in shallow sites and pelagic species like *S. microcephalus* and *P. glauca* in medium and deep sites (Figure 6). Species like *Labrus merula* and *Buenia affinis* were among the most abundant in number of reads (>1,000 per species) but have not been previously reported for the Bay of Biscay.

**Figure 5.**
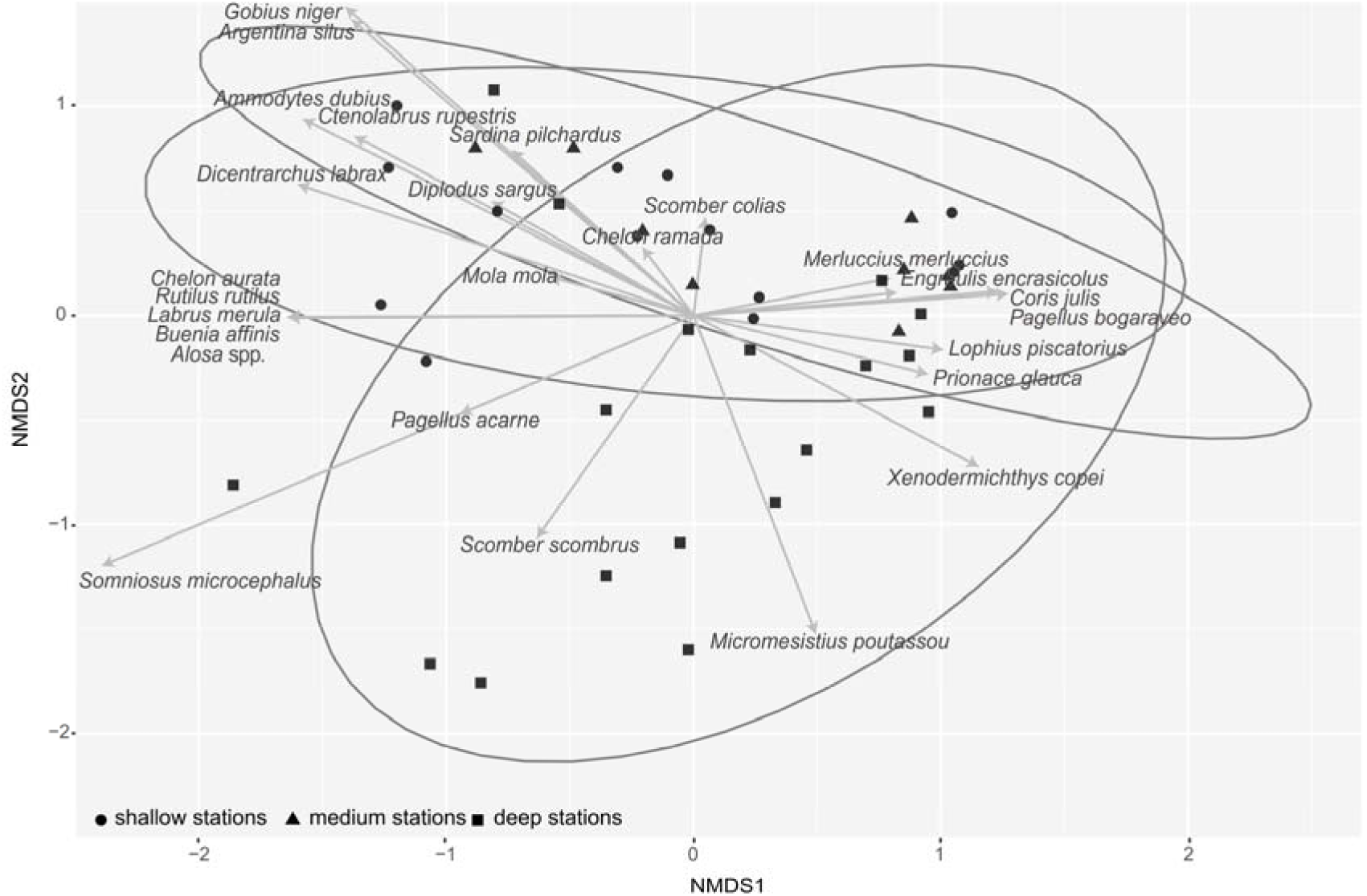
Non-metric multidimensional scaling (NMDS) plot, with a stress of 0.15, showing the similarity of species from each sample based on their relative abundance. The ellipse shows the 95% distance based on the centroid of the 3 sampling zones groups (coastal, neritic and oceanic). Spatial patterns of the species with >1,000 reads are shown.

**Figure 6.**
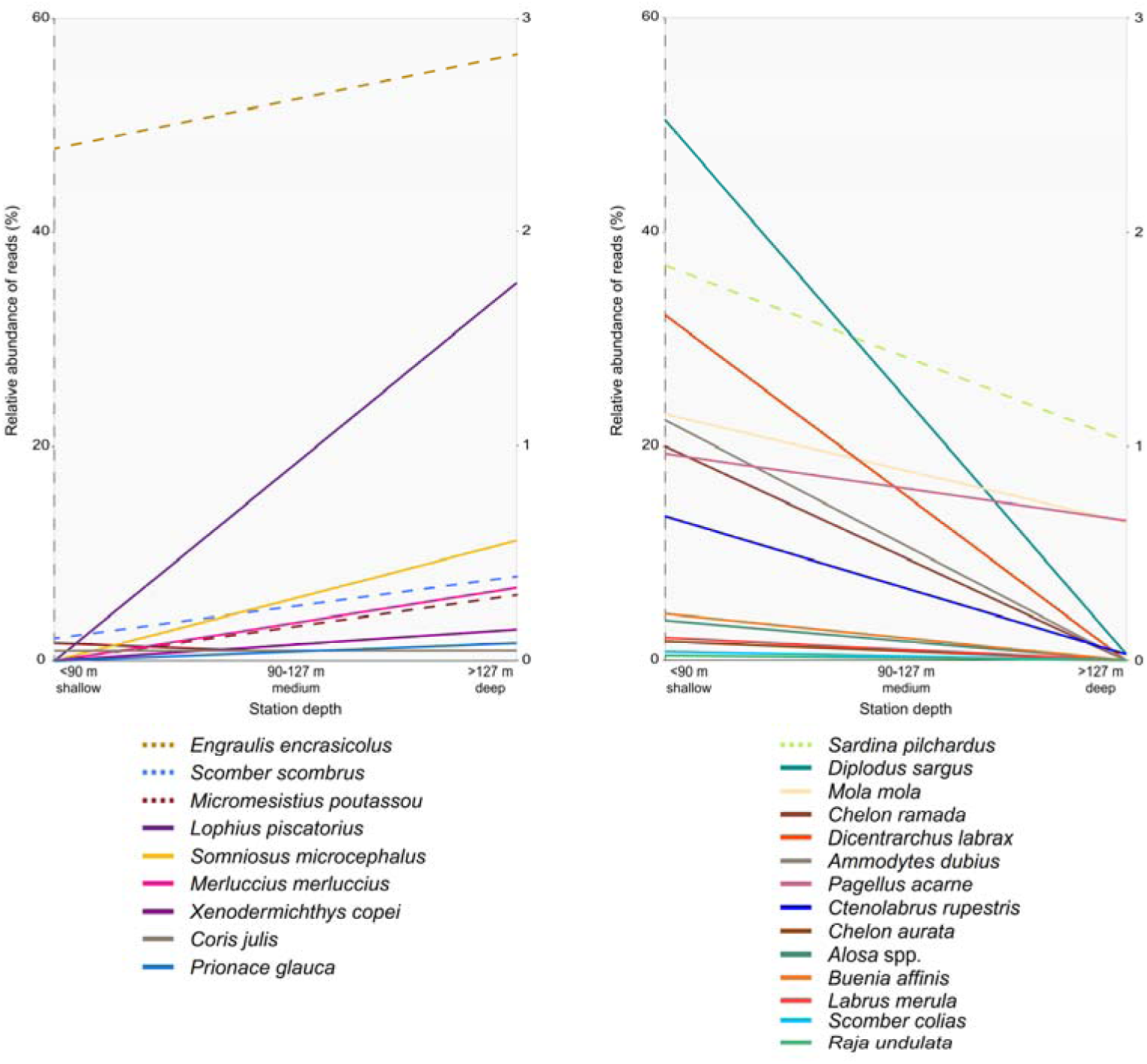
Linear relationship between depth and the relative number of reads obtained for each species with >1,000 reads. Scale of the right-hand y-axis has been amplified to ease visualization, dashed lines correspond to the left-hand y-axis, whilst continuous lines to the right-hand y-axis.

## Discussion

This study shows how eDNA metabarcoding provides a comprehensive overview of the fish diversity in a large-scale marine area. Compared to fish trawling, eDNA metabarcoding was able to “capture” a larger number of fish species. Both, eDNA and trawling based estimates (in number of reads and biomass, respectively) indicate that *E. encrasicolus* represents half of the abundance, which is consistent to the known large and stable anchovy population in the Bay of Biscay (Erauskin-Extramiana et al., 2019; Santos, Uriarte, Boyra, & Ibaibarriaga, 2018; Uriarte, Prouzet, & Villamor, 1996) and with the fact that the BIOMAN survey took place during the anchovy spawning season. The seven most abundant species in fish trawling representing >1% of the total biomass were *T. trachurus, S. scombrus, T. mediterraneus, M. merluccius, M. poutassou, S. pilchardus* and *B. boops*, which were all, except those not present in the reference database (*B. boops* and *T. mediterraneus*), also found in the eDNA metabarcoding data, and four of them (*E. encrasicolus, S. pilchardus*, *S. scombrus* and *M. poutassou*) were also among the most abundant species from eDNA data. Thus, concerning the most abundant species in the Bay of Biscay, eDNA and trawling data provided comparable conclusions.

The following three species were caught during fish trawling but were absent from eDNA data despite being present in the reference database, *Z. faber*, *S. sprattus* and *C. aper*. One possible explanation for this false negative detection could be the little abundance of this species’ DNA in the water, as suggested by the small and reduced number of catches (2.07 Kg in 3 sites, 1.07 Kg in 2 sites and 0.34 Kg in 2 sites, respectively). In fact, a small number of reads, i.e., 591, was also detected for *Solea solea*, a species from which 0.05 Kg were caught in a single station. If this is the case, filtering larger volumes of water and increasing sequencing depth could improve detection. Alternatively, reference sequences for *Z. faber S. sprattus* and *C. aper* could be undetected errors in the reference database (Li et al., 2018), or correspond to alternative intraspecific variants. On the other hand, in accordance with previous studies, eDNA data resulted in about 100 more species (35 with more than 10 reads) than trawling data collected simultaneously (Thomsen et al., 2012; Thomsen et al., 2016; Yamamoto et al., 2017). For example, species such as *D. sargus, P. acarne, M. mola, D. labrax, L. piscatorius, Chelon ramada, A. dubius, G. niger, Ctenolabrus ruperstris, A. silus, S. microcephalus and B. affinis* were not found in catches, but were more abundant in eDNA reads than the 5^th^ most abundant species (*M. merluccius*) in catches. The fact that eDNA results in a higher number of species could be partially attributed to the efficiency of the method to detect benthic or coastal species, difficult to catch by pelagic trawling nets, focused on small and medium-size pelagic species. To check to what extent eDNA is able to detect in surface waters (4 m) demersal species we compared the results with the ICES International Bottom Trawling Surveys (IBTS surveys) data for the Bay of Biscay from 2003 to 2019 (ICES, 2013) and with the 2017 Pélagiques Gascogne (PELGAS) integrated survey in the same area (Mathieu et al., 2019). eDNA metabarcoding data was able to detect at least 31 out of 164 species reported for the Bay of Biscay by IBTS surveys and 13 out of 45 species by PELGAS survey (Figure S5). Although not being a thorough comparison, as time periods and sampling seasons at least from IBTS surveys are different, the comparison provides an overall sense of eDNA as a potential method for surveying a large marine area in a relatively simple way. Differences in eDNA and pelagic trawl catchability can also explain the differences in relative abundances of the species found by the two kind of sampling methods, such as *S. pilchardus, M. poutassou* and *D. labrax*, with higher number of eDNA reads relative to the biomass caught, or *T. trachurus*, *S. scombrus* and *M. merluccius*, showing the opposite. However, similarity between both eDNA and trawling stations suggests that stations further apart tend to be more different. The amount, quality and stability of DNA molecules are largely affected by the production rate from each organism, diffusion of the molecules in the water and its inherent degradation (Collins et al., 2018; Murakami et al., 2019; Thomsen et al., 2012). But also, PCR amplification stochasticity and sequencing depth are known to affect the number of reads obtained from an eDNA sample (DiBattista et al., 2017; Zinger et al., 2019).

*T. minutus*, a morphologically similar species to *T. luscus* was identified through eDNA, which make us raise the hypothesis that specimens collected from catches were misidentified as *T. luscus*, potentially being *T. minutus* as eDNA revealed. This would not be an isolated case where morphological characteristics difficult to observe hamper taxonomic identification, and other available data (e.g., DNA) is needed for species identification (Dayrat, 2005). A remarkable case are lanternfishes of the Myctophidae, where species identification is based on the morphology and the shape and size of photophores, which are extremely fragile and seldom recovered intact (Cabrera-Gil et al., 2018). In this case, eDNA can play a major role for species identification as this study has shown, where at least five myctophid species were identified through eDNA. On the other hand, erroneous database records or missing sequences can bias eDNA based estimates. The quality and completeness of the reference database is crucial for taxonomic classification of eDNA data (Callahan, McMurdie, & Holmes, 2017). For example, two species were among the most abundant in our dataset, but not reported previously in the Bay of Biscay, namely *L. merula* and *B. affinis*. A careful examination suggests that, although *L. merula* could be misled by its close relative *L. bimaculatus*, occurring in the Bay of Biscay, the sequences attributed to *B. affinis* seem to be correctly assigned (Figure S6), suggesting that eDNA was able to detect species not previously reported in the area despite in low abundance.

Besides species diversity, eDNA also provides information on species distribution, which is comparable to that expected in the area. For instance, the number of reads assigned to the pelagic species *M. poutassou* and *S. scombrus* increased in stations deeper than 90m, where preferred habitats for these species occur (Ibaibarriaga et al., 2007). A contrasting pattern was observed for the greater argentine *A. silus*, a species commonly found at depths between 50 and 200m (Basterretxea et al., 2012), but found in our data at shallower stations. This could also suggest an incongruence with species identification with a close relative, in this case *A. sphyraena* commonly found over the continental slope (Basterretxea et al., 2012), but with no 12Sr RNA sequence in our reference database, or DNA from *A. silus* (even in its form of egg or larvae) dispersed to shallower stations. Similarly, species like *S. pilchardus, D. sargus, D. labrax, P. acarne* and *Alosa* spp. showed a distribution for this dataset in stations less than 90m depth, as our eDNA revealed. Available data on the diversity of elasmobranch species in the Bay of Biscay is limited, as most of these species are discarded from commercial fisheries and landing data is incomplete (ICES, 2017; Rodríguez-Cabello, Pérez, & Sánchez, 2013; Rusyaev & Orlov, 2013). Hence, in agreement to previous studies, our data supports eDNA as a potential mechanism for detecting and studying the distribution of elusive and deep-water species, which normally go undetected in fish trawl surveys, e.g., elasmobranchs, (Thomsen et al., 2016). In any case, eDNA results also revealed an ecological pattern for elasmobranchs. For instance *R. undulata*, which has a high-site fidelity occurred only in shallow waters (ICES, 2014), whilst large sharks *as S. microcephalus, P. glauca* and *Lamna nasus* predominantly occurred in deeper sites.

Aside from biological factors (e.g., individual shedding rate, persistence of DNA in the water) that can alter the quantity of eDNA released to the environment, technical considerations can introduce biases on the quality and number of reads generated per species and hence inferences driven from them (Dejean et al., 2011; Lamb et al., 2019; Thomsen et al., 2016). Reference databases are crucial to secure taxonomic assignment for data derived from eDNA samples (Zinger et al., 2019). While recent analyses on the taxonomic annotation of metazoan GenBank sequences suggest their reliability for eDNA metabarcoding studies (Leray, Knowlton, Ho, Nguyen, & Machida, 2019; Li et al., 2018), we encountered the need of including a thorough curation step for our ‘global’ database giving several mislabelled sequences. Species-level annotations were not considered in Leray et al. (2019), and we found incorrectly annotated sequences at all taxonomic levels. As environmental samples contain highly complex DNA signal from various organisms, primer choice is critical for species-level identification (Collins et al., 2019). We found that for our samples, the eukaryote universal COI primers result in a very small proportion of reads assigned to Actinopterygii. This is due to the fact that the primers target a large number of taxonomic groups, so larger coverage is needed for producing robust data (Alberdi, Aizpurua, Gilbert, Bohmann, & Mahon, 2018; Corse et al., 2019; Gunther, Knebelsberger, Neumann, Laakmann, & Martinez Arbizu, 2018; Stat et al., 2017). The use of more specific primers in our study allowed the specific detection of both Actinopterygii and Elasmobranchii. (Kelly, Port, Yamahara, & Crowder, 2014; Miya et al., 2015). Yet the amount of reads attributed to Elasmobranchii is small as ‘teleo’ primers were not specifically designed for this taxa, e.g., Kelly et al. (2014), and recent developments on elasmobranch-specific primers (Miya et al., 2015) could potentially be a powerful tool to increase the elasmobranch diversity in future marine surveys. In addition, for closely related species such as *Alosa alosa* and *Alosa fallax*, the target barcode was exactly the same, so being cautious we consider them as *Alosa* spp. Another crucial methodological step is the clustering method. We showed that using a clustering method (i.e., *vsearch* and *swarm*) decreased the number of identified species, probably because the algorithm merged sequences from different species into singular OTUs. Recent studies have suggested that clustering techniques and the use of percentages of similarities specially in short (<100 bp) sequences might mislead diversity estimates (Calderón-Sanou, Münkemüller, Boyer, Zinger, & Thuiller, 2019; Callahan et al., 2017; Xiong & Zhan, 2018). Thus, procuring a taxonomically comprehensive database with good quality sequences, and accurate data curation steps is crucial for producing robust and reproducible ecological conclusions from eDNA metabarcoding methods (Collins et al., 2019; Weigand et al., 2019). Including a human-specific blocking primer in our samples had little effect, as we indeed detect, although a small percentage (< 0.01%), reads identified as *H. sapiens*. The use of blocking primers in metabarcoding analysis has been previously used to block dominant taxa in a specific samples, for instance host DNA from diet analysis (Jakubavičiūtė, Bergström, Eklöf, Haenel, & Bourlat, 2017), or human DNA from ancient samples (Boessenkool et al., 2012). Our results suggest that our samples held very little contamination from external sources such as human manipulation, air or input from land.

Alternative ways to survey marine biodiversity and unbiased evaluations of the ecosystem components are needed as these provide the baseline for policy implementation in the context of global marine directives (e.g., Common Fisheries Policy or the Marine Strategy Framework Directive). eDNA metabarcoding is becoming a more accessible method that generates reliable information for ecosystem surveillance and invites its application on regular marine monitoring programs (Bohmann et al., 2014; Lacoursière-Roussel, Rosabal, & Bernatchez, 2016; Takahara, Minamoto, Yamanaka, Doi, & Kawabata, 2012). Results in this study have shown that eDNA samples provide information on fish diversity in a broad-scale marine area such as the Bay of Biscay. We detected almost ten times more fish and elasmobranch species compared with pelagic trawling. Some of these species can be considered elusive or difficult to capture with traditional fishing methods. In addition, ecological data was driven from eDNA, thus opening new avenues using molecular methods in broad-scale marine surveys.

## Acknowledgements

Authors are grateful to the crew of R/V Ramon Margalef and R/V Emma Bardán for their support during filtering and collection of samples, and specially to Luis Ferrer, Marina Chifflet, Bea Beldarrain and Carlota Pérez for their support on filtering onboard. Thanks are due to Iker Pereda for bioinformatic support and to Elisabete Bilbao for technical assistance. This project has been supported by the Department of Economic Development and Infrastructure of Basque Government (projects GENPES and ECOPES) and by the Spanish Ministry of Science, Innovation and Universities (project CTM2017-89500-R). This is contribution number X [to be included upon acceptance] from the Marine Research Division (AZTI).

