## Supplementary Figures for "Marine water environmental DNA metabarcoding provides a comprehensive fish diversity assessment and reveals spatial patterns in a large oceanic area"

^1^AZTI, Marine Research Division, Sukarrieta, Bizkaia, Spain.

^2^IKERBASQUE, Basque Foundation for Science, Bilbao, Spain.

^*^Corresponding author.

**Supplementary Figures**

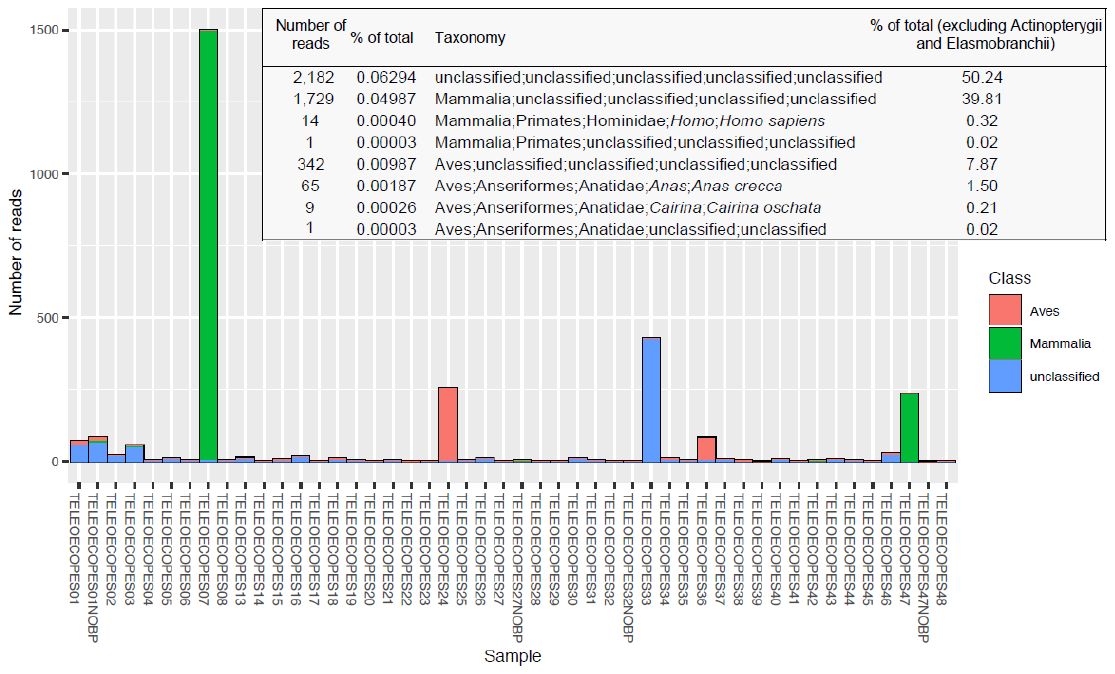

**Figure S1.** Taxonomic information of reads not classified as either Actinopterygii or Elasmobranchii when using the ‘global’ Chordata database and distribution of them in each of the eDNA samples.

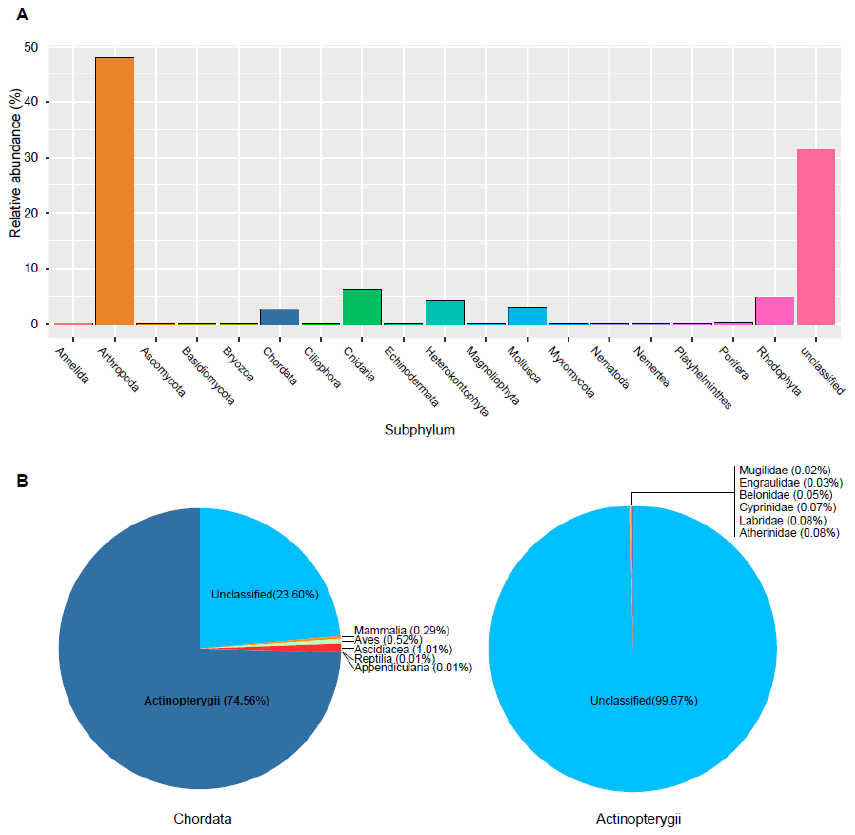

**Figure S2. (A)** Relative read abundance (%) of taxa classified to Subphylum, and **(B)** specifically classes within Chordata and families within Actinopterygii, respectively, from the four samples sequenced with the ‘mlCOI’ primers.

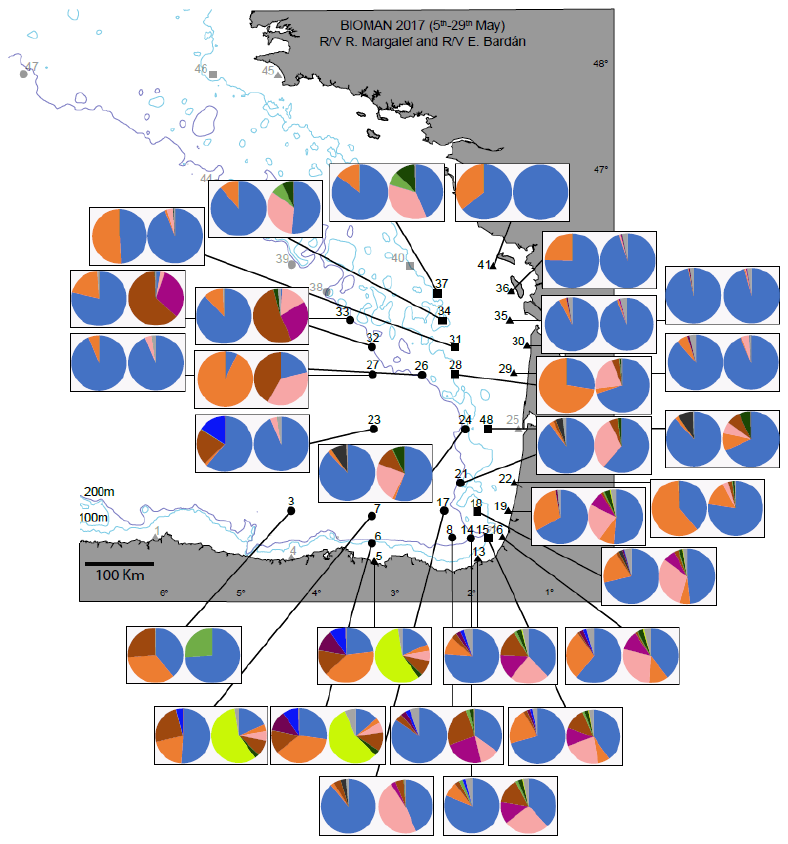

**Figure S3.** Pie charts showing the relative abundance of eDNA reads (first chart) and fish biomass caught (second chart) obtained from the 30 groups of stations within a 20nm ratio. eDNA charts include species with >10 reads only. Species with >5% biomass caught/number of reads per station are coded by colours, the rest are grouped in “others”.

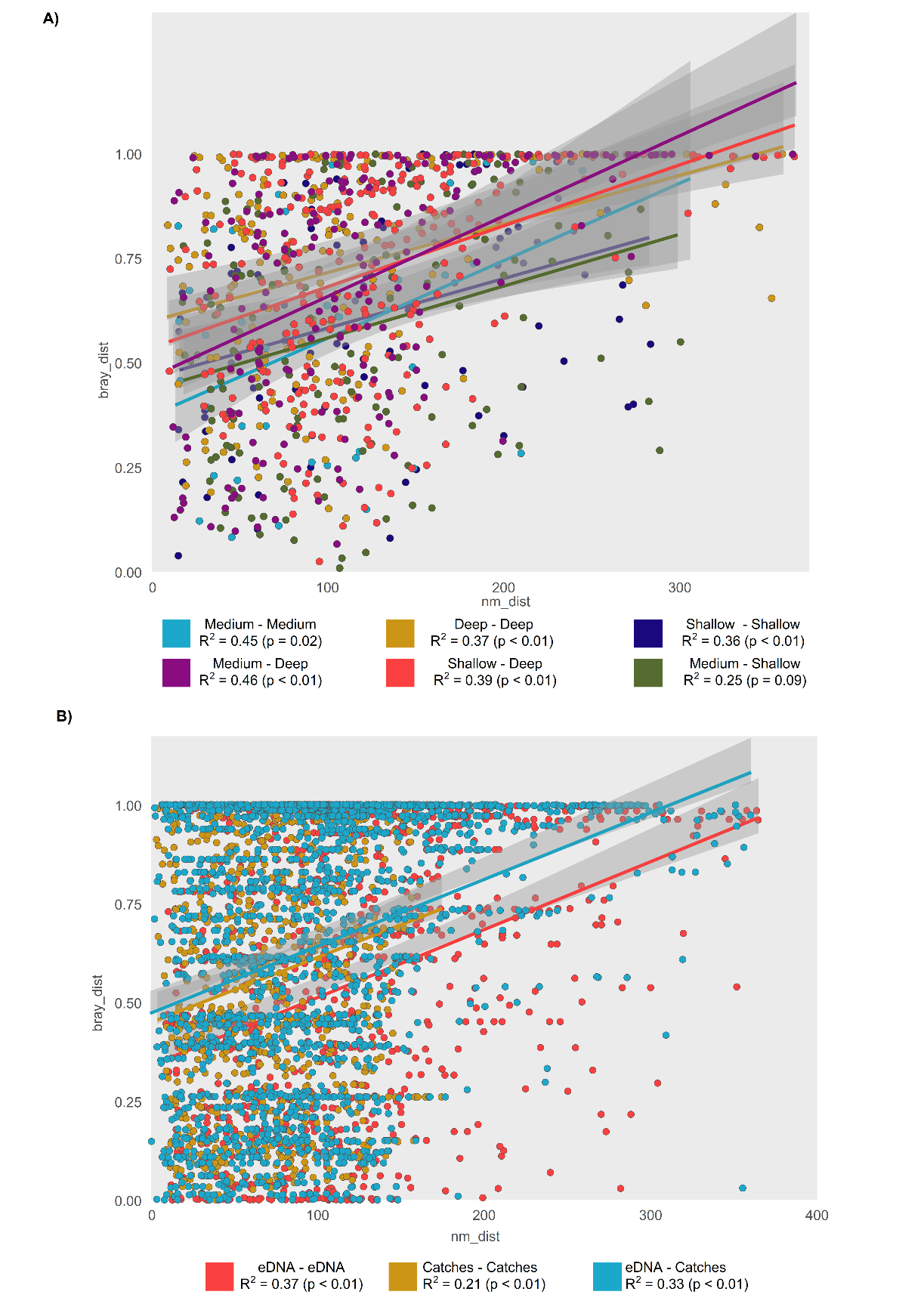

**Figure S4.** Scatterplot showing the relationship between Bray-Curtis distance and geographic distance between pairs of sampling points for **A)** eDNA, **B)** trawling and **C)** eDNA and trawling stations combined. Species included in C are only the common species detected by the two sampling methods. Pearson correlation is shown for each data group. Shaded area represents the 95% confidence interval of the linear regression.

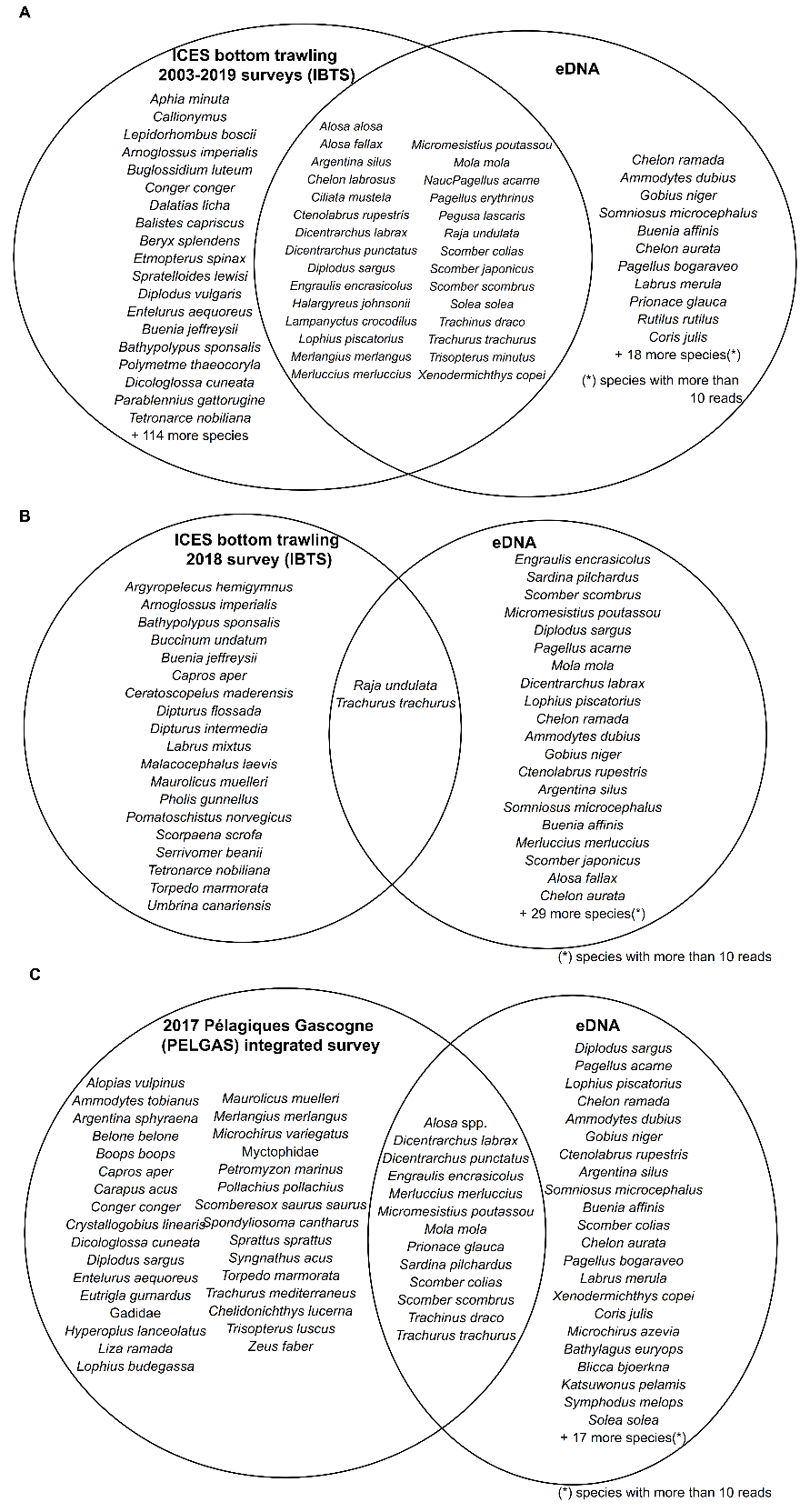

**Figure S5.** Venn diagrams showing fish caught in the ICES Bottom Trawling Survey carried out **A)** between 2003 and 2019 and **B)** in October 2018 available from ices.dk/marine-data/data-portals/ and **C**) in the 2017 Pélagiques Gascogne (PELGAS) integrated survey compared to the fish species detected through eDNA metabarcoding.

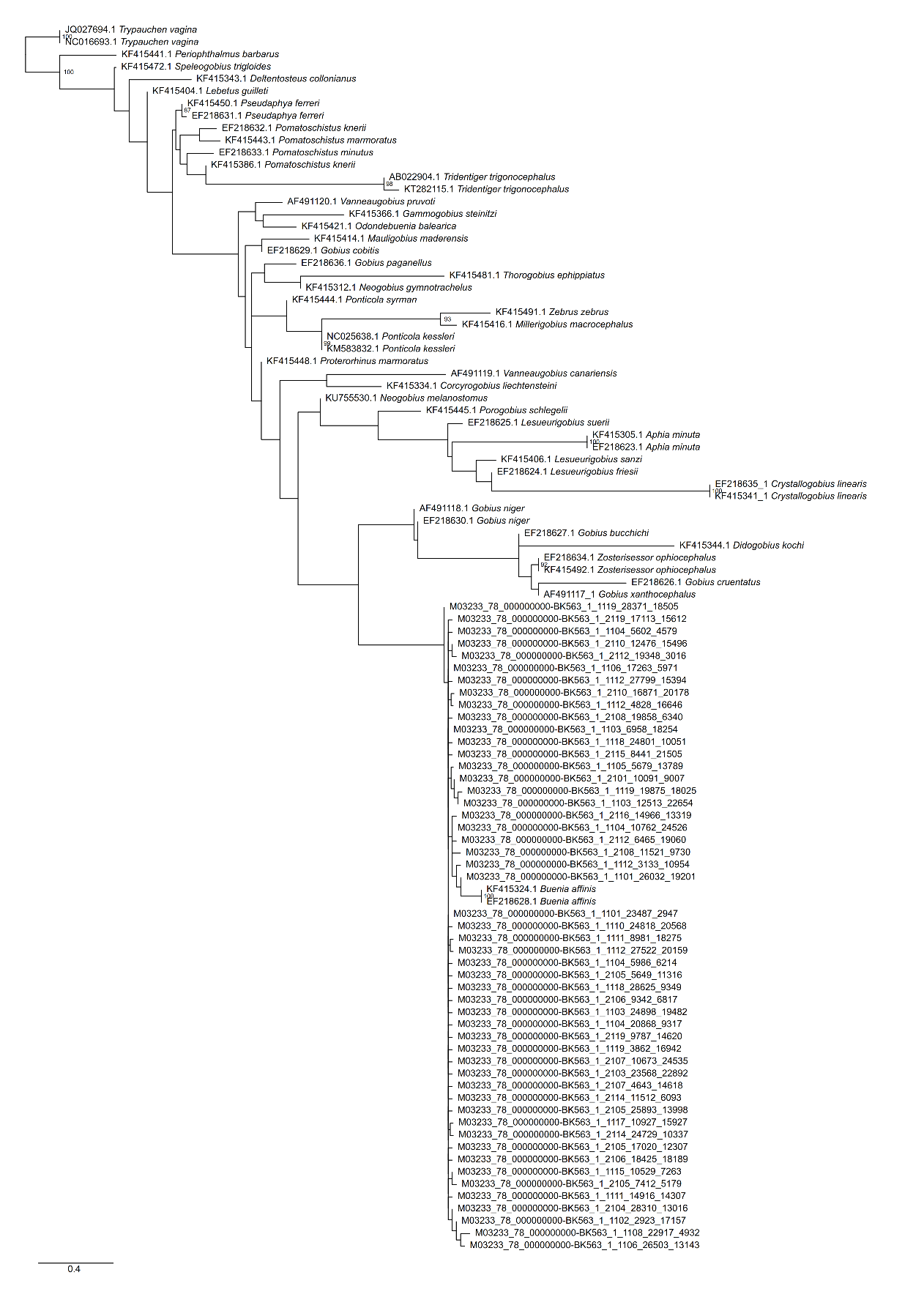

**Figure S6.** Maximum likelihood phylogenetic tree for species of Gobiidae included in the reference database and sequences obtained from eDNA samples putatively assigned to *Buenia affinis*. Bootstrap support values < 80% are not shown. Branch lengths indicate number of inferred substitutions per site. Phylogenetic tree constructed under the GTR+I+G model with 100 bootstrap replicates as implemented in RaxML (Stamatakis, 2014).
